# Differences in the transcriptome response in the gills of sea lamprey acutely exposed to 3-trifluoromethyl-4-nitrophenol (TFM), niclosamide or a TFM:niclosamide mixture

**DOI:** 10.1101/2023.06.08.544220

**Authors:** M.J. Lawrence, P. Grayson, J.D. Jeffrey, M.F. Docker, C.J. Garroway, J.M. Wilson, R.G. Manzon, M.P. Wilkie, K.M. Jeffries

## Abstract

Sea lamprey (*Petromyzon marinus*) control in the Laurentian Great Lakes of North America makes use of two pesticides: 3-trifluoromethyl-4-nitrophenol (TFM) and niclosamide, which are often co-applied. Sea lamprey appear to be vulnerable to these agents resulting from a lack of detoxification responses with evidence suggesting that lampricide mixtures produce a synergistic effect. However, there is a lack of information pertaining to the physiological responses of sea lamprey to niclosamide and TFM:niclosamide mixtures. Here, we characterized the transcriptomic responses of the sea lamprey to TFM, niclosamide, and a TFM:niclosamide (1.5%) mixture in the gill. Along with a control, larval sea lamprey were exposed to each treatment for 6 h, after which gill tissues were extracted for measuring whole-transcriptome responses using RNA sequencing. Differential gene expression patterns were summarized, which included identifying the broad roles of genes and common expression patterns among the treatments. While niclosamide treatment resulted in no differentially expressed genes, TFM- and mixture-treated fish had several differentially expressed genes that were associated with the cell cycle, DNA damage, metabolism, immune function, and detoxification. However, there was no common differential expression among treatments. For the first time, we characterized the transcriptomic response of sea lamprey to niclosamide and a TFM:niclosamide mixture and identified that these agents impact mRNA transcript abundance of genes associated with the cell cycle and cellular death, and immune function, which are likely mediated through mitochondrial dysregulation. These results may help to inform the production of more targeted and effective lampricides in sea lamprey control efforts.

## Introduction

The sea lamprey (*Petromyzon marinus*), native to the Atlantic Ocean, is an invasive species in the Laurentian Great Lakes of North America. It was documented in Lake Ontario in the 1880s and dispersed, via the Welland Canal, into Lake Erie and the upper Great Lakes in the early 1900s, resulting in severe ecological and economic damage due to parasitism on lake trout (*Salvelinus namaycush*) and other Great Lakes fishes (Smith and Tibbles 1980; Siefkes 2017; Wilkie et al. 2022). By the 1960s, sea lamprey control efforts made use of chemical pesticides applied to rivers and streams that are used as lampricides, targeting sea lamprey during their ∼4– 7-year larval stage burrowed in the sediment (Siefkes 2017; Wilkie et al. 2019). Two lampricides are used, 3-trifluoromethyl-4-nitrophenol (TFM), which primarily targets lampreys due to their relatively low capacity to detoxify TFM relative to other fishes (Lech and Statham 1975; Kane et al. 1993; Lawrence et al. 2021; Lawrence et al. 2022), and niclosamide, which is often mixed in small amounts (1-2 %) with TFM to increase the effectiveness of lampricide treatments without increasing toxicity to non-target fishes and other organisms (Wilkie et al. 2019). The greater toxicity of TFM-niclosamide mixtures results from synergistic or greater than additive interactions between the two compounds (e.g., Hepditch et al., 2021; Lawrence et al., 2021; Marking and Dawson, 1975), but how these compounds interact to enhance toxicity at the cellular level is relatively unknown. Consequently, a mechanistic understanding of the cellular effects of these agents is needed to better understand these interactions and to improve comprehension of how sea lamprey and other organisms respond to lampricides.

While these two lampricides have been the central focus of the sea lamprey control program, there is a scarcity of information on the physiological effects of acute lampricide exposures in a mixture on sea lamprey. The mode of action of TFM is through the uncoupling of oxidative phosphorylation in the mitochondria (Wilkie et al. 2007; Birceanu et al. 2009, 2011, 2014; Clifford et al. 2012; Lawrence et al. 2021), which decreases aerobic ATP production and forces the organism to rely upon anaerobic ATP synthesis, which is unsustainable over long periods. Gene ontogeny (GO) term enrichment analyses of RNA-sequencing (RNAseq) data have shown that acute TFM exposures are associated with transcriptome responses consistent with impairments to cellular growth and proliferation and enhanced apoptosis in bluegill and sea lamprey (Yin et al. 2021; Lawrence et al. 2022). The mechanism of action of niclosamide is thought to be through the uncoupling of oxidative phosphorylation in the mitochondria (Borowiec et al. 2022). This mode of action has been observed in both invertebrates (Weinbach and Garbus 1969; Jones 1979; Pampori et al. 1984; Pearson and Hewlett 1985) and fishes (Lawrence et al., 2021; Shoman, 2001; Zhu et al., 2020; Ionescu et al. 2022), which is often typified by depletion of tissue ATP and glycogen reserves, and increased production of lactate. In clinical studies, niclosamide has been identified as having effects on cell signalling pathways (e.g., wnt/β-catenin, mTOR, and STAT3) and promoting apoptosis, and its potential use for anti-cancer applications has been suggested (reviewed in Chen et al., 2018). More recently, niclosamide has been shown to have anti-viral properties in studies aimed at treating COVID-19 (Jeon et al. 2020; Weiss et al. 2021; Wotring et al. 2022). However, the effects of niclosamide in the transcriptome of fishes is mixed. In bluegill (*Lepomis macrochirus*), a teleost fish that is particularly lampricide-tolerant, exposure to niclosamide (0.068 mg L^-1^, a typical field dose used in lampricide applications) resulted in no differential expression of genes related to cellular growth and proliferation (Lawrence et al. 2023). In embryonic zebrafish (*Danio rerio*), niclosamide resulted in an enrichment of gene ontology (GO) terms associated with metabolism and cell cycle processes (Vliet et al. 2018; Zhu et al. 2020) with anatomical malformations occurring in juveniles (Zhu et al. 2019). Collectively, these findings suggest that niclosamide toxicity can have a mixed effect in fishes and can be associated with impaired energy metabolism and cellular growth processes.

Our understanding of the toxicology of TFM:niclosamide mixtures in fishes is limited. In bluegill exposed to a TFM:niclosamide mixture, there was an enrichment of GO terms associated with cellular growth, proliferation, and death observed (Lawrence et al. 2023). Furthermore, differential expression of several gene targets under TFM:niclosamide mixture treatment were associated with cellular growth signalling pathways (e.g., *wnt9b, notch1*, and *stat3*; Lawrence et al. 2023). Mixed lampricide exposures in fishes are also associated with a depletion of energy metabolites (e.g., glycogen, glucose, ATP, phosphocreatine) and lactate accumulation in tissues (Lawrence et al. 2021). In sea lamprey specifically, these effects were much more pronounced in the mixture treatment when compared to niclosamide or TFM alone, suggesting a synergistic mode of action (Lawrence et al. 2021).

This work aimed to address the molecular responses of acute exposures to niclosamide alone, TFM alone, and a TFM:niclosamide mixture in sea lamprey. Data for TFM alone are from a previous study on transcriptome responses to TFM in sea lamprey (Lawrence et al. 2022). We were interested in identifying expression patterns of genes involved in an exposure to TFM, niclosamide, and a mixture of the two to develop a better mechanistic understanding of the interaction between these two lampricides. In testing this, sea lamprey larvae were exposed to TFM alone (2.21 mg L^-1^ nominal), niclosamide alone (0.033 mg L^-1^ nominal), or a TFM:niclosamide (1.5%) mixture (TFM = 2.21 mg L^-1^, niclosamide = 0.033 mg L^-1^ nominal), for 6 h (see Lawrence et al. 2021 for details). The exposures represented the animal’s 24 h LC_10_ for TFM with 1.5 % of that as the nominal niclosamide concentration allowing us to characterize sublethal response pathways (Lawrence et al. 2021). Total RNA from gill tissue was extracted and sequenced using an RNAseq approach to characterize differential expression in the lamprey genome. We mainly focused on the gills of sea lamprey, which are in direct contact with the water and the route of lampricide uptake of lampreys, and therefore the first line of defence against these and other xenobiotics.

## Methods

This project was part of a larger study investigating the effects of TFM, niclosamide, and TFM:niclosamide mixtures on bluegill and sea lamprey transcriptomic responses. All projects used the same exposures, sampling, and analytical approaches, detailed elsewhere (Lawrence et al. 2021; Lawrence et al. 2022). Briefly, sea lamprey larvae (n = 568; TL = 104.94 ± 0.69 mm; mass = 1.47 ± 0.03 g) were obtained from tributaries of Lake Huron through electrofishing (Hammond Bay Biological Station, Millersburg, MI, USA) and were transported to Wilfrid Laurier University, Waterloo, Ontario, Canada, where they were held in flow-through circular tanks with an 8–10 cm deep layer of sand to promote natural burrowing behaviour (Lawrence et al. 2021). Experimental series were performed under the guidelines established by the Canadian Council of Animal Care with approval from the Wilfrid Laurier University Animal Care Committee (Animal Use Protocol No. R18001).

Following acclimation, lamprey were exposed to one of three experimental treatments for a total of 24 h: a control (dechlorinated tap water), niclosamide (0.033 mg L^-1^ nominal; measured 0.0224 ± 0.0012 mg L^-1^), TFM alone (2.21 mg L^-1^ nominal,; measured = 2.18 ± 0.02 mg L^-1^), or a TFM:niclosamide mixture (2.21 mg L^-1^ TFM, 0.033 mg L^-1^ niclosamide nominal; measured TFM = 2.54 ± 0.07 mg L^-1^, measured niclosamide = 0.0197 ± 0.0012 mg L^-1^). TFM concentrations were at the animals’ 24 h LC_10_ (i.e., TFM alone) as determined in preliminary toxicity trials on the same cohort of animals, and niclosamide values were set at 1.5% of the TFM concentration (see Lawrence et al. 2021). Tissue sampling of gill and liver occurred at 6, 12, and 24 h of exposure to develop a time course of effects. Animals were quickly euthanized using buffered tricaine methanesulfonate (MS-222; Syndel, Nanaimo, BC, Canada; 1.5 g L^-1^ MS-222 with 3.0 g L^-1^ NaHCO_3_), with gill and liver tissue being excised. Tissues were then preserved in RNAlater (Invitrogen, ThermoFisher Scientific, Mississauga, ON, Canada) and then stored at -80°C.

Total RNA from tissues was extracted using a commercial kit (RNeasy Plus Mini Kit; Qiagen, Toronto, ON, Canada). Quality control checks, including 260/280 and 260/230 absorbance ratios (NanoDrop One Microvolume UV-Vis SpectrophotometerThermoFisher, Mississauga, ON, Canada) and a Qubit RNA IQ assay (ThermoFisher, Mississauga, ON, Canada), were made prior to shipment to the sequencing facility. Sequencing and library preparation were performed by the Centre d’Expertise et de Services Génome Québec (Montreal, QC, Canada) on a NovaSeq 6000 sequencing system (Illumina, Vancouver, BC, Canada). Specific details on transcriptome assembly, alignment, and annotation can be found in Lawrence et al. (2022). Briefly, *de novo* transcriptomes were developed for sea lamprey with assemblies generated using Trinity (v2.8.5; Haas et al. 2013). Annotation of transcriptomes were handled using a Trinotate (v3.2) annotation protocol (http://trinotate.github.io) with SuperTranscripts being made via a Corset-Lace pipeline (https://github.com/Oshlack/Lace). While an annotated sea lamprey genome exists, we opted to use a *de novo* assembly as this project was making direct comparisons to bluegill, a species that has not been annotated to date (see Lawrence et al. 2022 for details). As this study represents a subset of that work, we used these same transcriptomes so that studies are comparable. Analysis of transcriptomes were performed in R studio (v1.3.1093) using the R programming language (v3.5.1; R Core Team 2020) with the package ‘edgeR’ (v3.24.3; Robinson et al. 2010; McCarthy et al. 2012) for filtering and differential expression analyses. For all analyses, complete time courses were performed for differential expression analyses (i.e., 6, 12, and 24 h). However, due to issues with high mortality in the mixture-exposed sea lamprey after 9 h (see Lawrence et al. 2021 for details) and resulting small sample sizes, we compared only 6 h exposed fish for all treatment groups. For clarity, differential expression in this study refers to differences in gene expression between a given lampricide treatment and the control (i.e., TFM vs control, niclosamide vs control, TFM:niclosamide mixture vs control).

The primary goals of this project were to test for common, differential gene expression patterns in the gills of TFM-, niclosamide-, and TFM:niclosamide mixture-exposed fish and to establish whether lampricide-responsive genes were consistent among our lampricide treatments. More specifically, we were interested in teasing apart how the TFM differential gene expression patterns change when fish are exposed to a TFM:niclosamide mixture. In doing so, we first screened for all differentially expressed clusters from a single control-treatment group comparison (e.g., gills of TFM vs control exposed fish). These differentially expressed clusters were then selected and compared to their response in the remaining two control-treatment comparisons regardless of whether they exhibited statistically significant differential expression through the whole-transcriptome analysis. This process was conducted for TFM and mixture-exposed fish but was not used for niclosamide-treated fish as there were no differentially expressed genes in the niclosamide only exposure. Patterns of common gene expression among the treatment groups (i.e., statistically significant differential expression, FDR < 0.05) were noted. Following this analysis, annotated clusters resulting from each treatment’s differentially expressed gene list were isolated and used in our gene role analyses as detailed below.

Identification of the annotated lamprey gene clusters for their broad role analysis was conducted using the information provided on UniProt (https://www.uniprot.org/). Here, each gene associated with a cluster was manually checked to determine which broad role group it belonged to using both the ‘function’ description on the gene’s entry page and the corresponding gene ontogeny (GO) terms. Criteria for each of the role groups are as follows. ‘Energy metabolism’ was associated with metabolic processes involved in mitochondrial function or enzymes/transporters involved in energy substrate translocation/processing. ‘Cell cycle/growth’ pertained to genes regulating a cell’s cycle function, cellular growth and proliferation, or apoptotic processes. ‘Immune’ corresponded to genes that regulate the immune functions, leukocyte functions, or the inflammatory response. ‘Ubiquitination’ involved genes regulating ubiquitination or that facilitated targeting during ubiquitination. ‘Detoxification’ included genes whose main function was eliminating xenobiotics. ‘DNA damage’ was associated with genes responsible for directly repairing DNA damage or those involved in the signalling of damage. ‘Nuclear process’ includes processes related to nuclear functions such as DNA replication, histone functions, and nuclear signalling. ‘RNA process’ involved those related to mRNA processing and functioning. ‘Receptor’ and ‘Transporter’ includes all genes that function principally as a receptor or a transporter, respectively, and are not related to any of the aforementioned processes. ‘Other’ gene functions refer to genes that were annotated with a function but did not fit into any of the above categories. There were also a few genes that had no corresponding role information through UniProt and have been annotated as ‘No info’ on the plots.

## Results

In the gills of TFM-exposed lamprey, downregulated superTranscript clusters were slightly more prevalent than upregulated ones, with 80 clusters and 59 clusters for downregulated and upregulated transcripts, respectively (Fig. 1). In contrast, mixture-exposed lamprey had a much larger number of upregulated (N = 60) transcripts than downregulated (N = 6) ones (Fig. 1) and had an overall lower number of differentially expressed transcripts when compared to TFM-exposed fish (66 total transcripts in mixture fish, 139 total transcripts in TFM fish). There was no differential expression associated with niclosamide treatment in the gills of sea lamprey.

**Figure 1:**
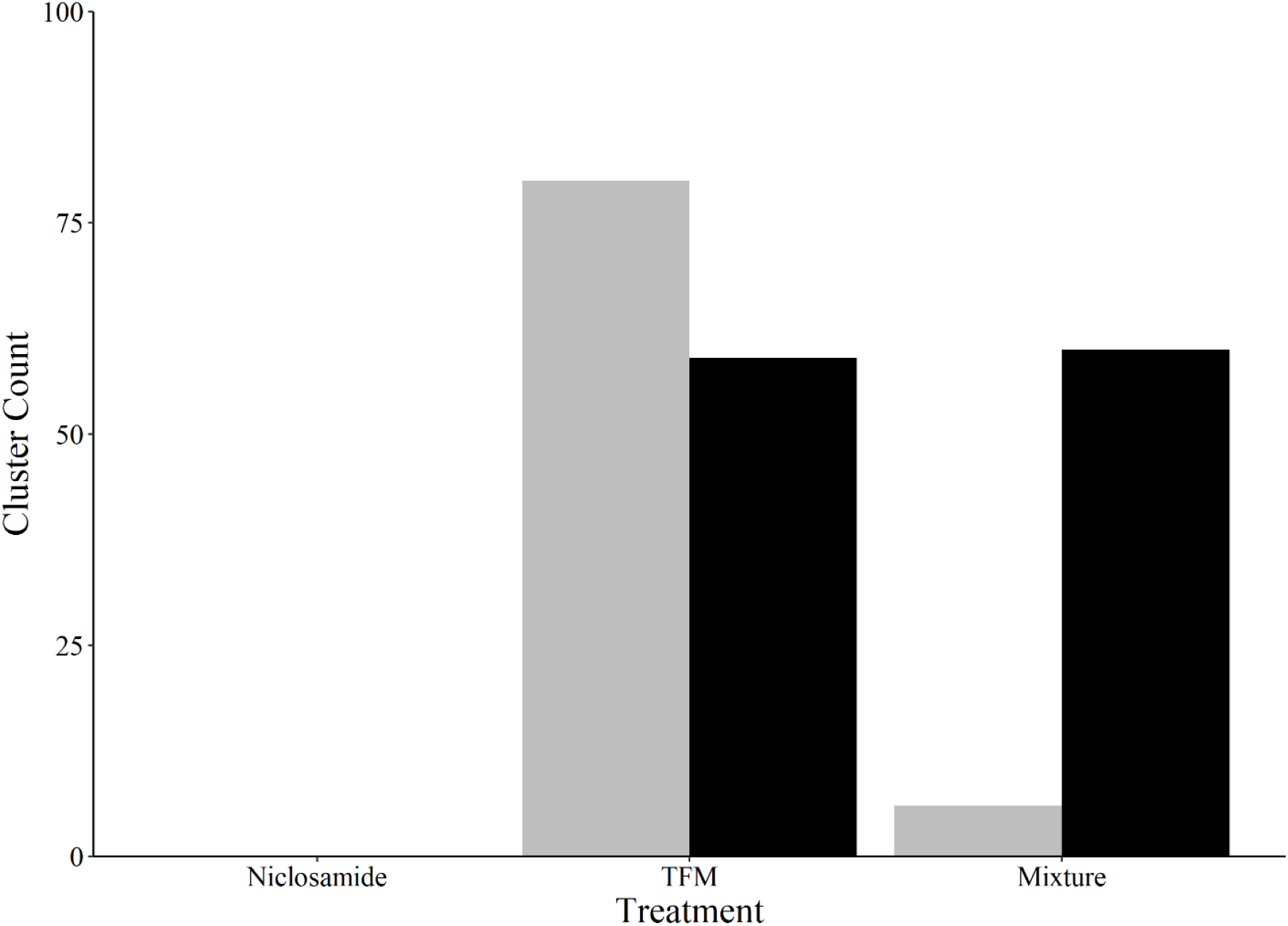
Total counts of differentially expressed superTranscriptome clusters in the gills of sea lamprey larvae at 6 h exposure to either niclosamide (0.033 mg L^-1^ nominal), TFM (2.21 mg L^-1^ nominal) or a TFM:niclosamide mixture (TFM = 2.21 mg L^-1^ nominal, niclosamide = 0.033 mg L^-1^ nominal). Liver data is not presented as only a single transcript was differentially expressed across all treatment groups, specifically *trib2* in mixture-treated fish. Grey bars represent differentially expressed clusters that were downregulated whereas black bars represent upregulated clusters. All clusters were deemed to be significant at a false discovery rate (FDR) of α = 0.05.

### Common mRNA transcript abundance patterns

One of the main objectives of this study was to characterize whether there were common patterns of gene expression among the three lampricide treatments. In doing so, we characterized transcripts that were differentially expressed in one treatment group and determined if these same clusters were differentially expressed in the other two treatments. However, of those transcripts which were differentially expressed in gills of TFM exposed lamprey relative to the controls (139 total), none of these were significantly differentially expressed in the gills of mixture or niclosamide-treated fish relative to the TFM treatment (Fig. 2A).

**Figure 2:**
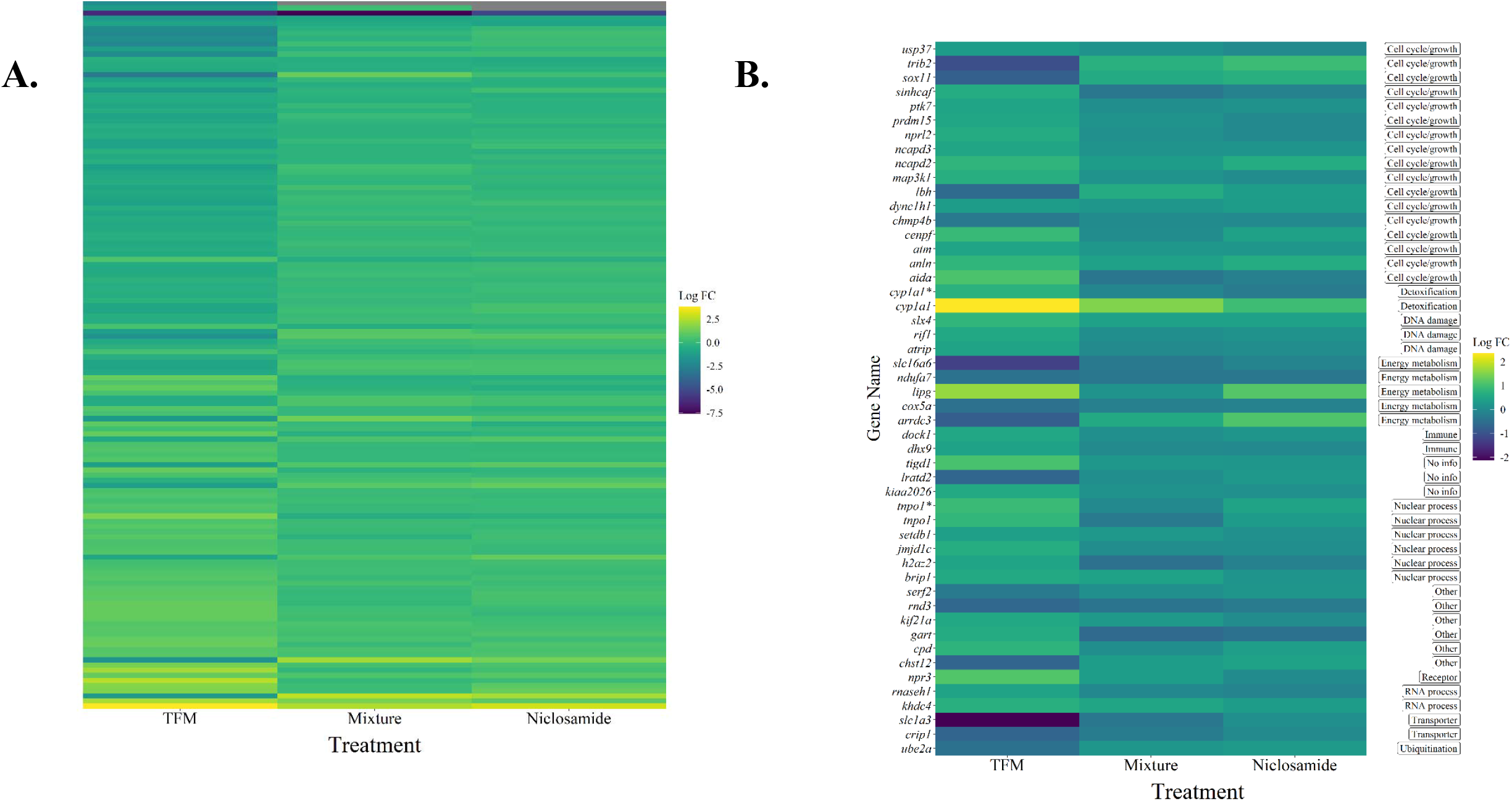
Heatmap depicting log fold changes (log FC) of all superTranscriptome clusters (**A**) and annotated clusters (**B**) that were differentially expressed in gills of larval sea lamprey exposed to TFM (2.21 mg L^-1^ nominal) and the responses of those same clusters in lamprey exposed to either a TFM:niclosamide mixture (TFM = 2.21 mg L^-1^ nominal, niclosamide = 0.033 mg L^-1^ nominal) or niclosamide (niclosamide = 0.033 mg L^-1^ nominal). Significant clusters in TFM exposed fish were deemed to be significant at a false discovery rate (FDR) of α = 0.05. In the niclosamide and mixture exposures, changes in gene expression were not significant (α > 0.05). In the case of annotated genes, the broad functional role of each gene can be found on the right side of the y-axis. Genes that associated with multiple clusters are delineated by an ‘*’ symbol.

In the mixture treatment, a total of 66 clusters were differentially expressed in the gills of sea lamprey with 25 of these clusters having annotations associated with them (Fig. 3). Despite all 66 of these clusters being found in both niclosamide- and TFM-treated fish, none of them were significantly differentially expressed in these groups (Fig. 3A). Collectively, our study showed that there were no common transcriptome patterns in the gills among TFM-, niclosamide-, and TFM:niclosamide mixture-treated sea lamprey after 6 h of exposure.

**Figure 3:**
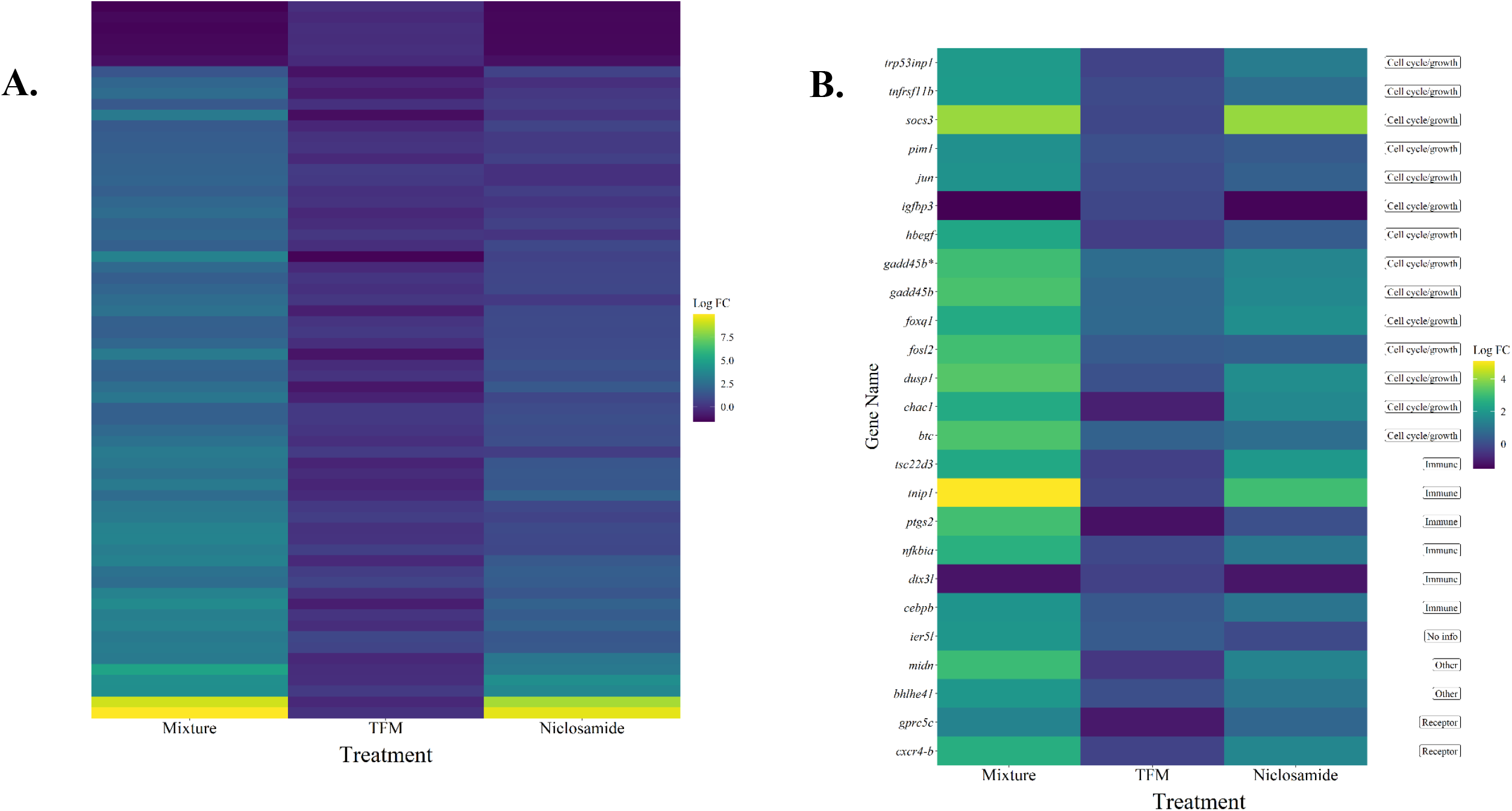
Heatmap depicting log fold changes (log FC) of all superTranscriptome clusters (**A**) and annotated clusters (**B**) that were differentially expressed in the gills of larval sea lamprey exposed to a TFM:niclosamide mixture (TFM = 2.21 mg L^-1^ nominal, niclosamide = 0.033 mg L^-1^ nominal) and the responses of those same clusters in lamprey exposed to either TFM (2.21 mg L^-1^ nominal) or niclosamide (niclosamide = 0.033 mg L^-1^ nominal). Significant clusters in mixture exposed fish were deemed to be significant at a false discovery rate (FDR) of α = 0.05. In the niclosamide and TFM exposures, the changes in gene expression were not significant (α > 0.05). In the case of annotated genes, the broad functional role of each gene can be found on the right side of the Y axis. Genes that associated with multiple clusters are delineated by an ‘*’ symbol.

### Gene analysis

In annotated clusters that were differentially expressed in the gills of TFM-treated fish, there were several patterns of interest from a functional perspective. Here, ‘cell cycle/growth’ constituted the largest role group consisting of mostly upregulated transcripts. Many of the genes in this group appeared to be involved with arresting cell cycle, growth, and proliferation (Fig. 2B). For example, TFM-exposure led to an upregulation of serine-protein kinase ATM (*atm*), which is believed to function in sensing DNA damage and helps to arrest the cell cycle and promote apoptosis. Similarly, a component of the GATOR1 complex, *nprl2*, was upregulated under TFM-treatment. This complex antagonizes mTORC1 signalling resulting in reduced cellular growth. Simultaneously, there was a downregulation of cellular proliferation and differentiation promoting factors such as *trib2* and *lbh*.

Exposure to TFM also resulted in an upregulation of several immune genes in the gill (Fig. 2B). This included an upregulation of ATP-dependent RNA helicase A (*dhx9*), which has a wide variety of functions related to transcriptional control of various components of the innate immune system, and dedicator of cytokinesis protein 1 (*dock1*) was also upregulated and is believed to play a role in facilitating clearance of apoptotic cells via phagocytosis. On a similar note, there was evidence for TFM-induced DNA damage as two genes involved in DNA damage sensing and repair mechanisms, *slx4* and *atrip*, were both upregulated. As might be expected, TFM exposures also corresponded with upregulation of genes associated with detoxification (*cyp1a1*) as well as affecting several genes involved with energy metabolism (Fig. 2B). In the latter case, this included genes associated with the electron transport chain in the mitochondria such as cytochrome c oxidase subunit 5A (*cox5a*) and NADH dehydrogenase 1 alpha subcomplex subunit 7 (*ndufa7*), both of which were downregulated in response to TFM (Fig. 2B). Interestingly, this corresponded with an upregulation of a lipase, *lipg*, which may function in triglyceride hydrolysis.

In contrast to the TFM-treatment, there was a lower diversity of role categories represented in the gills of mixture-exposed lamprey (Fig. 3B). Gene expression largely consisted of genes associating with ‘Cell cycle/growth’ and ‘Immune’ functions, with the former making up the largest portion of the role categories. While not sharing specific genes with TFM-exposed fish, there were some similar patterns exhibited in the gills of mixture-treated sea lamprey with respect to the ‘Cell cycle/growth’ role category. Specifically, there appeared to be upregulation of processes related to an arrest of cell cycle, proliferation, and growth, and apoptosis. For example, there was upregulation of *tnfrsf11b, chac1*, and *trp53inp1*, which are implicated in being pro-apoptotic factors (see Uniprot.org). Together, this suggests repressed growth and high cell death under exposure to the mixture treatment. Interestingly, some growth promoting genes including *btc* and *pim1* were upregulated under the mixture treatment, which demonstrates that the transcriptome response to TFM:nicolosamide treatment is complex.

As in the TFM-exposed lamprey, there was a large immune component in our differentially expressed clusters in mixture treated fish. This was largely related to pro-inflammatory processes, which were upregulated under this treatment. Interestingly, the only downregulated immune gene, *dtx3l*, was associated with DNA damage repair and interferons. Unlike TFM-exposed fish, there were no clusters associated with detoxification genes or energy metabolism in the gills of mixture treated fish.

## Discussion

### TFM:niclosamide mixture toxicity and transcriptome patterns

Sea lamprey exposed to the TFM:niclosamide mixture experienced transcriptomic changes associated with cell proliferation and immune function. While the specific physiological mechanisms underpinning mixed lampricide toxicity is poorly understood in fishes, this effect was consistent with what has been observed previously in teleosts (Lawrence et al. 2023). On its own, niclosamide is a potent regulator of cell cycle function and enhances cell death processes in a clinical setting (Jin et al. 2010; Hamdoun et al. 2017; Chen et al. 2018). Likewise, analogues of TFM (i.e., 2,4-dinitrophenol) have also been implicated in impairing cellular growth and increasing oxidative stress in clinical studies (Miyoshi et al. 2006; Han et al. 2008a, 2008b). In fishes, both TFM (Lawrence et al. 2022) and niclosamide (Vliet et al. 2018; Zhu et al. 2020) alone have been reported to show altered expression of genes associated with growth in fishes.

In fishes, changes in immune function and cellular growth/proliferation are believed to stem from the effect of these lampricides on mitochondrial respiration (Lam et al. 2013; Lawrence et al. 2022). Both TFM (Niblett and Ballantyne 1976; Birceanu et al. 2011; Huerta et al. 2020; Borowiec et al. 2022) and niclosamide (Tao et al. 2014; Kumar et al. 2018; Borowiec et al. 2022) have been identified as being potent uncouplers of mitochondrial respiration, with niclosamide being much more (40-60 fold) potent in this respect in sea lamprey (Borowiec et al. 2022). Increased rates of uncoupling could result in reactive oxygen species (ROS) generation (Nickel et al. 2014), which could result in repressed cellular growth, increased cellular death, and general inflammation responses (Lam et al. 2013). Our results support this as there was an upregulation of pro-apoptotic factors (e.g., *tnfrsf11b* and *chac1*), with a corresponding upregulation of growth repressing genes (e.g., *socs3*) in mixture-exposed sea lamprey.

Furthermore, a large component of the differentially expressed genes in the mixture treatment were associated with inflammatory responses, which further supports an involvement of mitochondrially derived ROS (López-Armada et al. 2013). While mixed lampricide toxicity in fishes has been shown to cause a depletion in energy substrates in sea lamprey (Lawrence et al. 2021), to our knowledge, this is the first work to show that mixed lampricide toxicity in sea lamprey includes a transcriptome response associated with effects on cellular proliferation and apoptosis.

Interestingly, exposure to the mixture treatment did not result in an altered expression of detoxification genes in sea lamprey after 6 h of exposure. Indeed, even in TFM alone, expression was limited to cytochrome p450’s (*cyp1a1*). As detailed below, this may be explained by a delayed transcriptional response in detoxification gene expression coupled with a reliance on sufficiently expressed or less plastic (inducible) detoxification proteins (i.e., steady state quantities of detoxification proteins; see *Niclosamide toxicity and transcriptome patterns*).

However, it is important to realize that by 12 h of exposure, most sea lamprey had died under the mixture treatment due to the relatively limited ability to detoxify TFM as previously observed (Lech 1974; Lech and Statham 1975; Kane et al. 1993; Bussy et al. 2018). Our recent analysis indicated that the genetic basis for the sea lamprey’s greater TFM sensitivity was due to a lower diversity of differentially expressed detoxification transcripts when compared to the more TFM-tolerant fishes such as bluegill (Lawrence et al. 2022), which can survive 8-9 fold higher concentrations of TFM, and 5-6 times higher concentrations of TFM-niclosamide mixtures (1 % niclosamide) than sea lamprey (Boogaard et al. 2003; Wilkie et al. 2019). Furthermore, previous work has identified that while sea lamprey have the capacity for conjugating TFM to a certain extent, the capacity to do so through the main biotransformation enzyme, uridine 5’-diphospho-glucuronosyltransferase (Ugt), appears limited (Kane et al. 1994; Bussy et al. 2018b, 2018a). The lack of detoxification response in mixture-exposed lamprey contrasts the response of bluegill where there was an upregulation of *cyp1a1, cyp1b1, sult1st1*, and *ugt3* by 6 h of exposure to the same TFM:niclosamide mixture (Lawrence et al. 2023). There was also a generally muted overall transcriptional response in sea lamprey, with only a handful of differentially expressed genes in all three treatment groups (Fig. 1), further explaining their relatively poor ability to detoxify lampricides (Lawrence et al. 2022). Together, this reinforces the fact that the lack of a rapid detoxification response of sea lamprey is likely a contributing factor governing the sensitivity of this species to lampricides.

### Niclosamide toxicity and transcriptome patterns

Exposure to niclosamide alone was not associated with differential gene expression, relative to controls, in sea lamprey. This was unexpected given that prior work showed a depletion of high energy substrates and an accumulation of parent (active) niclosamide in the liver and muscle of these same lamprey (Lawrence et al. 2021) and that niclosamide is more effective in uncoupling mitochondrial respiration in sea lamprey compared to TFM (Borowiec et al. 2022). The lack of a transcriptomic response in our study contrasts with teleosts that often show large-scale changes in the transcriptome following acute niclosamide exposures, targeting pathways related to energy metabolism, cellular growth and death, DNA damage, and detoxification, among others (Vliet et al. 2018; Zhu et al. 2020; Lawrence et al. 2023).

We propose that the observed lack of response in the transcriptome of niclosamide exposed sea lamprey at 6 h may result from the low exposure dosage used in our study. The niclosamide concentrations used here are well below a lethal dose level (Dawson et al. 1977) but are reflective of a niclosamide concentration that is 1.5% of TFM’s 24 h LC_10_ allowing us to characterize sublethal responses to these agents (Lawrence et al. 2021). This ratio of TFM to niclosamide is typical in field lampricide treatments (Brege et al. 2003). In fishes, niclosamide detoxification is believed to be through phase II biotransformation processes, namely glucuronidation and sulfation via UDP-glucuronylsyltransferase (Ugt) and Sulfotransferases (Sult), respectively (Statham and Lech 1975; Hubert et al. 2005). While the role of Ugts in sea lamprey appear limited (Kane et al. 1994; Bussy et al. 2018; Lawrence et al. 2023), Sult may be mediating detoxification of organic compounds in sea lamprey (Bussy et al. 2018; Lawrence et al. 2022). Thus, we might expect there to be some degree of niclosamide detoxification at substrate concentrations below the maximum enzyme reaction rate (Vmax), which could preclude any need for a transcriptional response. As we observed niclosamide accumulation in the tissues by 6 h of exposure (Lawrence et al. 2021), we may speculate that changes in gene expression may be evident at later time points, when the concentration of niclosamide in the livers were approximately 30 % higher (Lawrence et al. 2021), at which time substrate (niclosamide) concentrations could be nearer or greater than Vmax.

### Common patterns of lampricide toxicity

Our results demonstrated no common gene-level responses underlying lampricide toxicity in sea lamprey between the various lampricide treatments. This result may be consistent with the notion that toxicosis, while having similar manifestations at higher orders of biological scale, can result from several underlying molecular/physiological processes (Smart and Hodgson 2018). For example, exposures of zebrafish to two related phthalate compounds (i.e., reproductive toxins) resulted in similar effects on the fish’s reproductive physiology (e.g., decreased estradiol levels, altered ovarian histology) but had different gene targets when observing the transcriptomic profiles, suggesting differences in the mechanism of action of these compounds (Chen et al. 2019). Similarly, a literature review of the effects of the phenylurea herbicides diuron and linuron in fishes indicated that were different gene targets between these agents (Marlatt and Martyniuk 2017). An alternative explanation may be that the lack of shared gene response may be a result of the mixture-exposed fish tending towards death as there was considerable mortality by 9 h of exposure (Lawrence et al. 2021). In this case, the affected genes may simply represent a status where the cellular function is in a much more dysregulated state in the mixture-exposed fish relative to TFM-exposed fish. This is especially evident in that mixture fish did experience mortality such that differences in transcriptome response may simply represent a signature of death or a more progressed toxicological state. Furthermore, there were fewer differentially expressed transcripts in the mixture-exposed fish, relative to TFM-exposed fish, with a shift towards a greater proportion of upregulated clusters suggesting an altered response to the mixture treatment. Indeed, this explanation is consistent with the much larger physiological perturbations seen in mixture-treated lamprey against all other groups (Lawrence et al. 2021). In support of this, previous works have identified dose-dependent transcriptome patterns in fishes exposed to varying concentrations of toxicants (Santos et al. 2010; Griffitt et al. 2013; Mittal et al. 2022), suggesting that gene expression patterns are reflective of the degree of toxicity. Indeed, the question of shared gene targets becomes increasingly complex when one considers the unknown actions that the interaction of TFM and niclosamide have when placed in a mixture. It is established that there is synergism between TFM and niclosamide at the physiological level (Hepditch et al. 2021; Lawrence et al. 2021), which could suggest that the unique gene targets in the mixture-treated fish observed here are in part responsible for mediating this effect. However, further enzyme kinetic and toxicological analyses are needed to learn more about the cellular underpinnings of the mixture interaction of lampricides.

### Lampricide toxicology and sea lamprey control efforts

The use of lampricides has been an integral part of sea lamprey control in the Great Lakes for decades (Wilkie et al. 2022). In the last few years, our fundamental understanding of the toxicology and species-specific processes underlying the more selective action of lampricides has improved (Bussy et al. 2018b, 2018a; Huerta et al. 2020; Birceanu et al. 2021; Lawrence et al. 2021, 2022; Borowiec et al. 2022). Indeed, transcriptomics has become increasingly important in understanding the mechanisms underlying lamprey sensitivity to lampricides and the holistic range of toxicological responses to these agents (Lawrence et al. 2021, 2022, 2023; Yin et al. 2021; this study). It appears that, in conjunction with mitochondria assessments (Borowiec et al. 2022), TFM and niclosamide toxicity is mediated through mitochondrial dysregulation that may result in ROS-induced damage in both lamprey and teleosts (Lawrence et al. 2022, 2023; this study). Importantly, sensitivity appears to result from the ability to rapidly eliminate lampricides from the body through phase II biotransformation, namely glucuronidation. In our earlier works, we established that the larger diversity and magnitude of changes in detoxification transcripts likely conferred a higher tolerance to TFM in bluegill, relative to sea lamprey (Lawrence et al. 2022). Furthermore, bluegill exposed to a TFM:niclosamide mixture showed differential expression of a diverse set of detoxification transcripts which were still observed in niclosamide-only treatments, albeit at lower levels of change than the mixture fish (Lawrence et al. 2023). This contrasts what was observed here in that all three lampricide treatments did not have a significant effect on detoxification gene expression, further reenforcing the idea that higher sensitivity in lamprey likely stems from a muted ability to detoxify lampricides.

This knowledge may aid in developing more effective and selective lampricides in sea lamprey controls efforts (Lantz et al. 2022). Based on our molecular assessments (Lawrence et al. 2021, 2022, 2023; Yin et al. 2021; this study), we believe that the general lack of glucuronidation via Ugt in sea lamprey may be exploited to develop more targeted lampricides. As Ugts are believed to have evolved in response to detoxifying plant-derived toxins (Bock 2016), it may present opportunities for using naturally derived and biodegradable plant toxins that are effective in targeting lamprey but are rather innocuous to other native fishes in the area. Similarly, synthetic compounds that are structurally similar to TFM/niclosamide may prove to be useful if their primary means of detoxification includes catalysis by Ugt. For example, Lam et al. (2013) found that 4-Nitrophenol, a compound structurally similar to TFM, produced many of the same molecular responses in zebrafish that were found with TFM exposures in bluegill and sea lamprey (Lawrence et al. 2022), suggesting that structural similarity may be important in deriving a new effective synthetic pesticide. While identification of specific compounds is beyond the scope of this work, our transcriptomic research provides a foundational framework by which these compounds could be identified using a primary screening of the mechanism of action or through the primary elimination pathways. The identification of novel lampricides is quickly becoming an important consideration in sea lamprey control efforts as there is concern that sea lamprey may develop resistance to these pesticides (Dunlop et al. 2018; Lantz et al.

2022). Transcriptomics is believed to be an important tool in the development of next generation lampricides (Lantz et al. 2022) and these data provide some of the first stages in identifying the exploitable pathways and systems for sea lamprey-specific lampricides.

## Acknowledgements

The authors would like to thank Oana Birceanu for logistical assistance, Fisheries and Oceans Canada for providing the TFM used in this study, Darren Foubister and Josh Sutherby for conducting the exposures, Kimberly Ta for mRNA extractions, and the crew at the U.S. Geological Survey’s Hammond Bay Biological Station for collecting the larval lamprey used in this study.

## Author Information

KMJ, MPW, RGM, JMW, CJG, and MFD were responsible for the experimental design. The labs of KMJ and MPW carried out the exposures and sample collection. Data and sample analyses were conducted by MJL, PG, and JDJ. The first draft of the manuscript was written by MJL and PG, with all authors contributing to the refinement and production of the final manuscript. MJL and PG contributed equally to this paper.

## References

Birceanu O., G.B. McClelland, Y.S. Wang, J.C.L. Brown, and M.P. Wilkie. 2011. The lampricide 3-trifluoromethyl-4-nitrophenol (TFM) uncouples mitochondrial oxidative phosphorylation in both sea lamprey (Petromyzon marinus) and TFM-tolerant rainbow trout (Oncorhynchus mykiss). Comparative Biochemistry and Physiology Part C: Toxicology & Pharmacology 153:342–349.

Birceanu O., G.B. McClelland, Y.S. Wang, and M.P. Wilkie. 2009. Failure of ATP supply to match ATP demand: The mechanism of toxicity of the lampricide, 3-trifluoromethyl-4-nitrophenol (TFM), used to control sea lamprey (Petromyzon marinus) populations in the Great Lakes. Aquatic Toxicology 94:265–274.

Birceanu O., L.A. Sorensen, M. Henry, G.B. McClelland, Y.S. Wang, and M.P. Wilkie. 2014. The effects of the lampricide 3-trifluoromethyl-4-nitrophenol (TFM) on fuel stores and ion balance in a non-target fish, the rainbow trout (Oncorhynchus mykiss). Comparative Biochemistry and Physiology Part C: Toxicology & Pharmacology 160:30–41.

Birceanu O., L.R. Tessier, B. Huerta, W. Li, A. McDonald, and M.P. Wilkie. 2021. At the intersection between toxicology and physiology: What we have learned about sea lampreys and bony fish physiology from studying the mode of action of lampricides. Journal of Great Lakes Research, Supplement on Sea Lamprey International Symposium III (SLIS III) 47:S673–S689.

Bock K.W. 2016. The UDP-glycosyltransferase (UGT) superfamily expressed in humans, insects and plants: Animal-plant arms-race and co-evolution. Biochemical Pharmacology 99:11–17.

Borowiec B.G., O. Birceanu, J.M. Wilson, A.E. McDonald, and M.P. Wilkie. 2022. Niclosamide Is a Much More Potent Toxicant of Mitochondrial Respiration than TFM in the Invasive Sea Lamprey (Petromyzon marinus). Environ Sci Technol.

Brege D.C., D.M. Davis, J.H. Genovese, T.C. McAuley, B.E. Stephens, and R. Wayne Westman. 2003. Factors Responsible for the Reduction in Quantity of the Lampricide, TFM, Applied Annually in Streams Tributary to the Great Lakes from 1979 to 1999. Journal of Great Lakes Research 29:500–509.

Bussy U., Y.-W. Chung-Davidson, T. Buchinger, K. Li, S.A. Smith, A. Daniel Jones, and W. Li. 2018a. Metabolism of a sea lamprey pesticide by fish liver enzymes part B: method development and application in quantification of TFM metabolites formed in vivo. Anal Bioanal Chem 410:1763–1774.

Bussy U., Y.-W. Chung-Davidson, T. Buchinger, K. Li, S.A. Smith, A.D. Jones, and W. Li. 2018b. Metabolism of a sea lamprey pesticide by fish liver enzymes part A: identification and synthesis of TFM metabolites. Anal Bioanal Chem 410:1749–1761.

Chen H., W. Feng, K. Chen, X. Qiu, H. Xu, G. Mao, T. Zhao, et al. 2019. Transcriptomic analysis reveals potential mechanisms of toxicity in a combined exposure to dibutyl phthalate and diisobutyl phthalate in zebrafish (Danio rerio) ovary. Aquatic Toxicology 216:105290.

Chen W., R.A. Mook, R.T. Premont, and J. Wang. 2018. Niclosamide: Beyond an antihelminthic drug. Cellular Signalling 41:89–96.

Clifford A.M., M. Henry, R. Bergstedt, D.G. McDonald, A.S. Smits, and M.P. Wilkie. 2012. Recovery of Larval Sea Lampreys from Short-Term Exposure to the Pesticide 3-Trifluoromethyl-4-Nitrophenol: Implications for Sea Lamprey Control in the Great Lakes. Transactions of the American Fisheries Society 141:1697–1710.

Dawson V.K., K.B. Cummings, and P.A. Gilderhus. 1977. Efficacy of 3-trifluoromethyl-4-nitrophenol (TFM), 2’, 5-dichloro-4’-nitrosalicylanilide (Bayer 73), and a 98: 2 Mixture of Lampricides in Laboratory Studies. US Fish and Wildlife Service.

Dunlop E.S., R. McLaughlin, J.V. Adams, M. Jones, O. Birceanu, M.R. Christie, L.A. Criger, et al. 2018. Rapid evolution meets invasive species control: the potential for pesticide resistance in sea lamprey. Can J Fish Aquat Sci 75:152–168.

Griffitt R.J., C.M. Lavelle, A.S. Kane, N.D. Denslow, and D.S. Barber. 2013. Chronic nanoparticulate silver exposure results in tissue accumulation and transcriptomic changes in zebrafish. Aquatic Toxicology 130–131:192–200.

Haas B.J., A. Papanicolaou, M. Yassour, M. Grabherr, P.D. Blood, J. Bowden, M.B. Couger, et al. 2013. De novo transcript sequence reconstruction from RNA-seq using the Trinity platform for reference generation and analysis. Nat Protoc 8:1494–1512.

Hamdoun S., P. Jung, and T. Efferth. 2017. Drug Repurposing of the Anthelmintic Niclosamide to Treat Multidrug-Resistant Leukemia. Front Pharmacol 8.

Han Y.H., S.W. Kim, S.H. Kim, S.Z. Kim, and W.H. Park. 2008a. 2,4-Dinitrophenol induces G1 phase arrest and apoptosis in human pulmonary adenocarcinoma Calu-6 cells. Toxicology in Vitro 22:659–670.

Han Y.H., S.Z. Kim, S.H. Kim, and W.H. Park. 2008b. 2,4-Dinitrophenol induces apoptosis in As4.1 juxtaglomerular cells through rapid depletion of GSH. Cell Biology International 32:1536–1545.

Hepditch S.L.J., O. Birceanu, and M.P. Wilkie. 2021. A Toxic Unit and Additive Index Approach to Understanding the Interactions of 2 Piscicides, 3LTrifluoromethylL4LNitrophenol and Niclosamide, in Rainbow Trout. Environ Toxicol Chem 40:1419–1430.

Hubert T.D., J.A. Bernardy, C. Vue, V.K. Dawson, M.A. Boogaard, T.M. Schreier, and W.H. Gingerich. 2005. Residues of the Lampricides 3-Trifluoromethyl-4-nitrophenol and Niclosamide in Muscle Tissue of Rainbow Trout. J Agric Food Chem 53:5342–5346.

Huerta B., Y.-W. Chung-Davidson, U. Bussy, Y. Zhang, J.N. Bazil, and W. Li. 2020. Sea lamprey cardiac mitochondrial bioenergetics after exposure to TFM and its metabolites. Aquatic Toxicology 219:105380.

Jeon S., M. Ko, J. Lee, I. Choi, S.Y. Byun, S. Park, D. Shum, et al. 2020. Identification of antiviral drug candidates against SARS-CoV-2 from FDA-approved drugs. Antimicrobial agents and chemotherapy 64:e00819–20.

Jin Y., Z. Lu, K. Ding, J. Li, X. Du, C. Chen, X. Sun, et al. 2010. Antineoplastic Mechanisms of Niclosamide in Acute Myelogenous Leukemia Stem Cells: Inactivation of the NF-κB Pathway and Generation of Reactive Oxygen Species. Cancer Res 70:2516–2527.

Jones W.E. 1979. Niclosamide as a Treatment for Hymenolepis Diminuta and Dipylidium Caninum Infection in Man *. The American Journal of Tropical Medicine and Hygiene 28:300–302.

Kane A.S., M.W. Kahng, R. Reimschuessel, P.T. Nhamburo, and M.M. Lipsky. 1994. UDP-Glucuronyltransferase Kinetics for 3-Trifluoromethyl-4-nitrophenol (TFM) in Fish. Transactions of the American Fisheries Society 123:217–222.

Kumar R., L. Coronel, B. Somalanka, A. Raju, O.A. Aning, O. An, Y.S. Ho, et al. 2018. Mitochondrial uncoupling reveals a novel therapeutic opportunity for p53-defective cancers. Nat Commun 9:3931.

Lam S.H., C.Y. Ung, M.M. Hlaing, J. Hu, Z.-H. Li, S. Mathavan, and Z. Gong. 2013. Molecular insights into 4-nitrophenol-induced hepatotoxicity in zebrafish: Transcriptomic, histological and targeted gene expression analyses. Biochimica et Biophysica Acta (BBA) - General Subjects 1830:4778–4789.

Lantz S.R., R.A. Adair, J.J. Amberg, R.A. Bergstedt, M.A. Boogaard, U. Bussy, M.F. Docker, et al. 2022. Next Generation Lampricides: A Three-Stage Process to Develop Improved Control Tools for Invasive Sea Lamprey. Canadian Journal of Fisheries and Aquatic Sciences 79:692–702.

Lawrence M.J., P. Grayson, J.D. Jeffrey, M.F. Docker, C.J. Garroway, J.M. Wilson, R.G. Manzon, et al. 2022. Variation in the Transcriptome Response and Detoxification Gene Diversity Drives Pesticide Tolerance in Fishes. Environ Sci Technol 56:12137–12147.

Lawrence M.J., P. Grayson, J.D. Jeffrey, M.F. Docker, C.J. Garroway, J.M. Wilson, R.G. Manzon. 2023. Transcriptomic impacts and potential routes of detoxification in a lampricide-tolerant teleost exposed to TFM and niclosamide. Comparative Biochemistry and Physiology Part D: Genomics and Proteomics 46:101074.

Lawrence M.J., D. Mitrovic, D. Foubister, L.M. Bragg, J. Sutherby, M.F. Docker, M.R. Servos, et al. 2021. Contrasting physiological responses between invasive sea lamprey and non-target bluegill in response to acute lampricide exposure. Aquatic Toxicology 237:105848.

López-Armada M.J., R.R. Riveiro-Naveira, C. Vaamonde-García, and M.N. Valcárcel-Ares. 2013. Mitochondrial dysfunction and the inflammatory response. Mitochondrion 13:106–118.

Marking L.L. and V.K. Dawson. 1975. Method for assessment of toxicity or efficacy of mixtures of chemicals (Report No. 67). Investigations in Fish Control.

Marlatt V.L. and C.J. Martyniuk. 2017. Biological responses to phenylurea herbicides in fish and amphibians: New directions for characterizing mechanisms of toxicity. Comparative Biochemistry and Physiology Part C: Toxicology & Pharmacology 194:9–21.

McCarthy D.J., Y. Chen, and G.K. Smyth. 2012. Differential expression analysis of multifactor RNA-Seq experiments with respect to biological variation. Nucleic Acids Research 40:4288–4297.

Mittal K., J. Ewald, and N. Basu. 2022. Transcriptomic Points of Departure Calculated from Rainbow Trout Gill, Liver, and Gut Cell Lines Exposed to Methylmercury and Fluoxetine. Environmental Toxicology and Chemistry 41:1982–1992.

Miyoshi N., H. Oubrahim, P.B. Chock, and E.R. Stadtman. 2006. Age-dependent cell death and the role of ATP in hydrogen peroxide-induced apoptosis and necrosis. Proceedings of the National Academy of Sciences 103:1727–1731.

Niblett P.D. and J.S. Ballantyne. 1976. Uncoupling of oxidative phosphorylation in rat liver mitochondria by the lamprey larvicide TFM (3-trifluoromethyl-4-nitrophenol). Pesticide Biochemistry and Physiology 6:363–366.

Nickel A., M. Kohlhaas, and C. Maack. 2014. Mitochondrial reactive oxygen species production and elimination. Journal of Molecular and Cellular Cardiology 73:26–33.

Pampori N.A., G. Singh, and V.M.L. Srivastava. 1984. Cotugnia digonopora: carbohydrate metabolism and effect of anthelmintics on immature worms*. Journal of Helminthology 58:39–47.

Pearson R.D. and E.L. Hewlett. 1985. Niclosamide Therapy for Tapeworm Infections. Ann Intern Med 102:550–551.

R Core Team (2020). — European Environment Agency. n.d. Methodology Reference.

Robinson M.D., D.J. McCarthy, and G.K. Smyth. 2010. edgeR: a Bioconductor package for differential expression analysis of digital gene expression data. Bioinformatics 26:139–140.

Santos E.M., J.S. Ball, T.D. Williams, H. Wu, F. Ortega, R. Van Aerle, I. Katsiadaki, et al. 2010. Identifying health impacts of exposure to copper using transcriptomics and metabolomics in a fish model. Environmental science & technology 44:820–826.

Shoman H. 2001. Ultrastructural changes associated with molluscicide hepatotoxicity in fishes. Egypt J of Aquatic Biolo and Fish 5:219–236.

Siefkes M.J. 2017. Use of physiological knowledge to control the invasive sea lamprey (Petromyzon marinus) in the Laurentian Great Lakes. Conservation Physiology 5:cox031.

Siefkes M.J., T.B. Steeves, W.P. Sullivan, M.B. Twohey, and W. Li. 2013. Sea Lamprey Control: Past, Present, and Future. Pp. 651–704 in W.W. Taylor, A.J. Lynch, and N.J. Leonard eds. Great Lakes Fisheries Policy and Management. Michigan State University Press, East Lansing, MI, USA.

Smart R.C. and E. Hodgson, eds. 2018. Molecular and Biochemical Toxicology (Fifth edition.). Wiley, Hoboken, NJ.

Smith B.R. and J.J. Tibbles. 1980. Sea Lamprey (Petromyzon marinus) in Lakes Huron, Michigan, and Superior: History of Invasion and Control, 1936–78. Can J Fish Aquat Sci 37:1780–1801.

Statham C.N. and J.J. Lech. 1975. Metabolism of 2′,5-Dichloro-4′-Nitrosalicylanilide (Bayer 73) in Rainbow Trout (Salmo gairdneri). J Fish Res Bd Can 32:515–522.

Tao H., Y. Zhang, X. Zeng, G.I. Shulman, and S. Jin. 2014. Niclosamide ethanolamine–induced mild mitochondrial uncoupling improves diabetic symptoms in mice. Nat Med 20:1263–1269.

Vliet S.M., S. Dasgupta, and D.C. Volz. 2018. Niclosamide Induces Epiboly Delay During Early Zebrafish Embryogenesis. Toxicological Sciences.

Weinbach E.C. and J. Garbus. 1969. Mechanism of Action of Reagents that uncouple Oxidative Phosphorylation. Nature 221:1016–1018.

Weiss A., F. Touret, C. Baronti, M. Gilles, B. Hoen, A. Nougairède, X. de Lamballerie, et al. 2021. Niclosamide shows strong antiviral activity in a human airway model of SARS-CoV-2 infection and a conserved potency against the Alpha (B. 1.1. 7), Beta (B. 1.351) and Delta variant (B. 1.617. 2). PloS one 16:e0260958.

Wilkie M.P., J.A. Holmes, and J.H. Youson. 2007. The lampricide 3-trifluoromethyl-4-nitrophenol (TFM) interferes with intermediary metabolism and glucose homeostasis, but not with ion balance, in larval sea lamprey (Petromyzon marinus). Can J Fish Aquat Sci 64:1174–1182.

Wilkie M.P., T.D. Hubert, M.A. Boogaard, and O. Birceanu. 2019. Control of invasive sea lampreys using the piscicides TFM and niclosamide: Toxicology, successes & future prospects. Aquatic Toxicology 211:235–252.

Wilkie M.P., N.S. Johnson, and M.F. Docker. 2022. Invasive species control and management: The sea lamprey story.

Wotring J.W., S.M. McCarty, K. Shafiq, C.J. Zhang, T. Nguyen, S.R. Meyer, R. Fursmidt, et al. 2022. In Vitro Evaluation and Mitigation of Niclosamide’s Liabilities as a COVID-19 Treatment. Vaccines 10:1284.

Yin X., A.S. Martinez, A. Perkins, M.M. Sparks, A.M. Harder, J.R. Willoughby, M.S. Sepúlveda, et al. 2021. Incipient resistance to an effective pesticide results from genetic adaptation and the canalization of gene expression. Evol Appl 14:847–859.

Zhu B., W. He, F. Yang, and L. Chen. 2020. High-throughput transcriptome sequencing reveals the developmental toxicity mechanisms of niclosamide in zebrafish embryo. Chemosphere 244:125468.

Zhu B., B. Li, Q. Feng, and Y. Xu. 2019. Neurotoxicity of niclosamide on juvenile zebrafish. Chinese Journal of Parasitology and Parasitic Diseases 37:588–592.

